# The House Fly Y Chromosome is Young and Minimally Differentiated from its Ancient X Chromosome Partner

**DOI:** 10.1101/073023

**Authors:** Richard P. Meisel, Christopher A. Gonzales, Hoang Luu

**Affiliations:** Department of Biology and Biochemistry, University of Houston

**Author notes:** Current address: Department of Pharmacology, UT Southwestern Medical Center. To whom correspondence should be addressed:Richard Meisel, 3455 Cullen Blvd #342, Houston, TX 77204-5001.

**Keywords:** sex chromosomes, neo-Y chromosome, *Musca domestica*

## Abstract

Canonical ancient sex chromosome pairs consist of a gene rich X (or Z) chromosome and a male- (or female-) limited Y (or W) chromosome that is gene poor. In contrast to highly differentiated sex chromosomes, nascent sex chromosome pairs are homomorphic or very similar in sequence content. Nascent sex chromosomes arise frequently over the course of evolution, as evidenced by differences in sex chromosomes between closely related species and sex chromosome polymorphisms within species. Sex chromosome turnover typically occurs when an existing sex chromosome becomes fused to an autosome or an autosome acquires a new sex-determining locus/allele. Previously documented sex chromosome transitions involve changes to both members of the sex chromosome pair (X and Y, or Z and W). The House fly has sex chromosomes that resemble the ancestral fly karyotype that originated 100 million years ago, and therefore house fly is expected to have X and Y chromosomes with different gene content. We tested this hypothesis using whole genome sequencing and transcriptomic data, and we discovered little evidence for genetic differentiation between the X and Y in house fly. We propose that house fly has retained the ancient X chromosome, but the ancestral Y was replaced by an X chromosome carrying a male determining gene. Our proposed hypothesis provides a mechanisms for how one member of a sex chromosome pair can experience evolutionary turnover while the other member remains unaffected.

## 1. Introduction

In organisms where sex is determined by genetic factors, sex determining loci can reside on sex chromosomes. Sex chromosome systems are divided into two broad categories: 1) males are the heterogametic sex (XY); or 2) females are the heterogametic sex (ZW). In long established sex chromosomes—such as in birds, eutherian mammals, and *Drosophila*—the X and Y (or Z and W) chromosomes are typically highly differentiated (Charlesworth, 1996; Charlesworth et al., 2005). The X (or Z) chromosome usually resembles an autosome in size and gene density, although there are some predicted and observed differences in gene content and evolutionary rates between the X (or Z) and autosomes (Rice, 1984; Charlesworth et al., 1987; Vicoso and Charlesworth, 2006; Sturgill et al., 2007; Ellegren, 2011; Meisel et al., 2012; Meisel and Connallon, 2013). In contrast, Y (Z) chromosomes tend to contain a small number of genes with male- (female-) specific functions and are often enriched with repetitive DNA as a result of male- (female-) specific selection pressures, a low recombination rate, and a reduced effective population size (Rice, 1996; Bachtrog, 2013). This X-Y (or Z-W) differentiation results in a heterogametic sex that is effectively haploid for most or all X (or Z) chromosome genes.

Highly divergent X-Y (or Z-W) pairs trace their ancestry to a pair of undifferentiated autosomes (Bull, 1983; Charlesworth, 1991). Many species harbor undifferentiated sex chromosomes because they are either of recent origin or non-canonical evolutionary trajectories have prevented X-Y (or Z-W) divergence (Stöck et al., 2011; Bachtrog, 2013; Vicoso et al., 2013; Yazdi and Ellegren, 2014). Recently derived sex chromosomes often result from Robertsonian fusions between an existing sex chromosome and an autosome, or they can arise through a mutation that creates a new sex determining locus on an autosome (Bachtrog et al., 2014; Beukeboom and Perrin, 2014). In both cases, one of the formerly autosomal homologs evolves into an X (or Z) chromosome, and the other homolog evolves into a Y (or W) chromosome. In some cases, one or both of the ancestral sex chromosomes can revert back to an autosome when a new chromosome becomes sex-linked (Carvalho and Clark, 2005; Larracuente et al., 2010; Vicoso and Bachtrog, 2013). In all of the scenarios described above, the X and Y (or Z and W) chromosomes evolve in concert, with an evolutionary transition in one sex chromosome producing a corresponding change in its partner.

Sex chromosome evolution has been extensively studied in higher dipteran flies (Brachycera), where sex chromosome transitions involving X-autosome fusions are common (Patterson and Stone, 1952; Schaeffer et al., 2008; Baker and Wilkinson, 2010; Vicoso and Bachtrog, 2015). The ancestral brachyceran karyotype consists of five large autosomal pairs (known as Muller elements A-E) and a heterochromatic, gene-poor sex chromosome pair (element F is the X chromosome); this genomic arrangement has been conserved for ~100 million years (my) in some lineages (Muller, 1940; Foster et al., 1981; Weller and Foster, 1993; Vicoso and Bachtrog, 2013). In species with the ancestral karyotype, females are XX and males are XY, with a male-determining locus (M factor) on the Y chromosome (Bopp et al., 2014). Many sex chromosome transitions have occurred across Brachycera, including fusions of ancestral autosomes with the X chromosome and complete reversions of the ancestral X to an autosome (Baker and Wilkinson, 2010; Vicoso and Bachtrog, 2013, 2015).

The house fly (*Musca domestica*) is a classic model system for studying sex determination because it harbors a vast array of natural and laboratory genetic variation (Dübendorfer et al., 2002). The house fly karyotype resembles that of the ancestral brachyceran, with five large euchromatic elements and a heterochromatic sex chromosome pair (Boyes et al., 1964). The house fly X and Y chromosomes can be distinguished based on their length in cytological preparations (Denholm et al., 1983; Cakir and Kence, 1996; Hediger et al., 1998b), and close relatives of the house fly (*Lucilia* blow flies) have the ancestral karyotype with differentiated X and Y chromosomes (Linger et al., 2015; Vicoso and Bachtrog, 2015). In contrast to other flies, the house fly M factor has been mapped to each of the five autosomes in addition to the Y chromosome (Hamm et al., 2015). The autosomes harboring M are expected to be neo-Y chromosomes, but the house fly Y chromosome is often assumed to be the ancestral brachyceran Y that is differentiated from the X (Vicoso and Bachtrog, 2013; Hamm et al., 2015). However, there are multiple reasons to suspect that the house fly Y chromosome is not differentiated from the X in terms of gene content. First, no sex-linked genetic markers have been identified on the ancestral House fly sex chromosomes other than M (Hamm et al., 2015), suggesting that there are no X-specific genes or genetic variants. Second, males with an autosomal M factor that do not carry a Y chromosome are fertile (Bull, 1983; Hamm et al., 2015), suggesting that no essential male fertility genes are unique to the Y chromosome apart from the M factor. Third, house flies that carry only a single copy of either the X or Y chromosome (i.e., XO or YO flies) are viable and fertile (Bull, 1983; Hediger et al., 1998a), indicating that no essential genes are uniquely found on the X and missing from the Y chromosome and vice versa.

We used whole genome and transcriptome sequencing to test for genetic differentiation between between the house fly X and Y chromosomes. We observed minimal differentiation in sequence and gene content between X and Y chromosomes. We propose that the ancestral Y chromosome has been lost from house fly populations, and that existing Y chromosomes in natural populations arose from the recent translocation of the M factor onto an ancestral X chromosome. This represents, to the best of our knowledge, the first example of the “recycling” of a sex chromosome pair through the creation of a nascent Y from an ancient X chromosome (Graves, 2005).

## 2. Results

### 2.1. The House fly X and Y chromosomes do not have unique sequences

Our first goal was to identify house fly X chromosome sequences not found on the Y, which would be consistent with the hypothesis that house flies have an ancient, differentiated sex chromosome pair. Males of the house fly genomic reference strain (aabys) have been previously characterized as possessing the XY karyotype (Wagoner, 1967; Tomita and Wada, 1989; Scott et al., 2014). To identify X-linked genes and examine differentiation between X and Y chromosomes, we used the Illumina technology to sequence genomic DNA (gDNA) separately from male (XY) and female (XX) aabys flies (3 replicates of each sex), and we aligned the reads to the annotated genome (read counts are available in **Supplemental Data S1**). If house fly males have a Y chromosome that is fully differentiated from the X, we expect females to have twice the sequencing coverage 
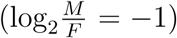
 within genes on Muller element F (the ancestral X chromosome) as males (Vicoso and Bachtrog, 2013). We instead find that the average sequencing coverage in males and females is almost identical 
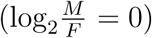
 for genes on all six chromosomes (Fig 1).

**Figure 1:**
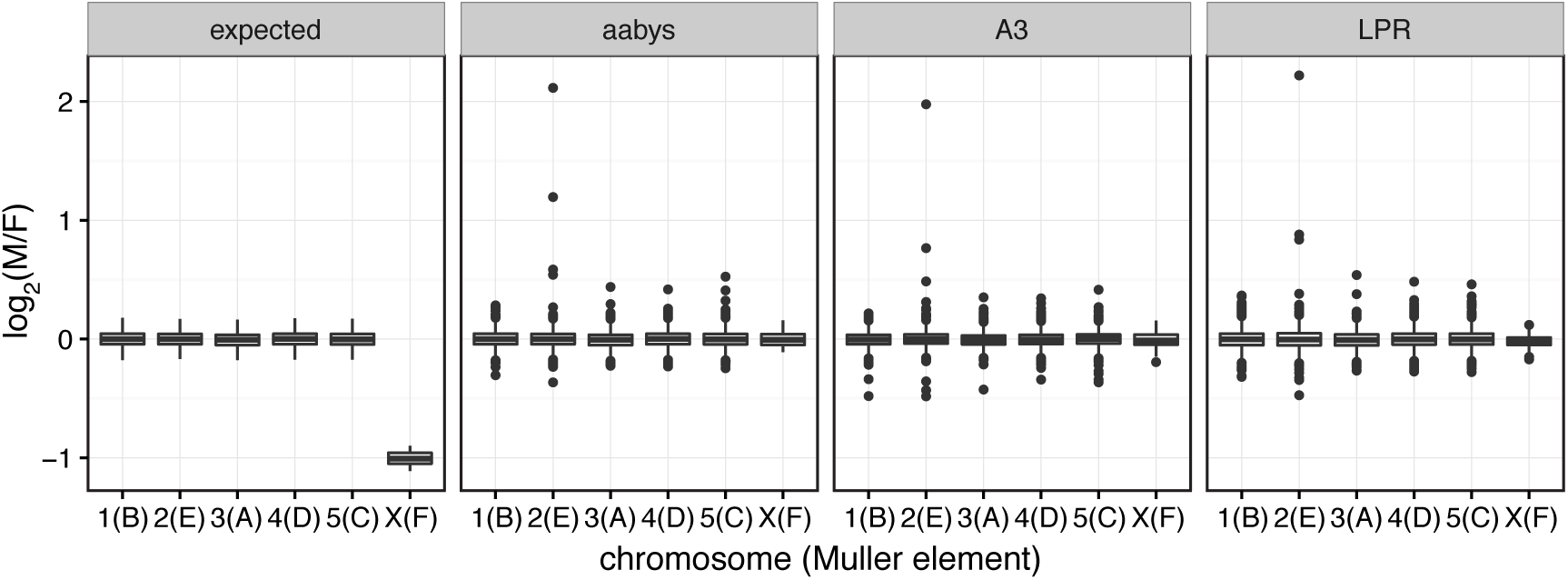
Expected sequencing coverage in males relative to females (
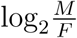
) in an XY system with a degenerated Y chromosomes (left), and observed coverage in three House fly strains (aabys, A3, and LPR) for each House fly chromosome (Muller elements in parentheses).

To determine whether lack of X-Y differentiation is common to other XY strains of the house fly, we sought to identify X-linked genes in two additional strains previously reported to have XY males: A3 and LPR (Scott and Georghiou, 1985; Scott et al., 1996; Liu and Yue, 2001). We sequenced gDNA from males and females of the A3 and LPR strains, and we aligned those reads to the reference genome (read counts are available in **Supplemental Data S2 & S3**). Consistent with the results from aabys, both the A3 and LPR strains had equal sequencing coverage within genes across all six chromosomes in males and females (**Fig 1**). In addition 
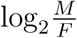
, is moderately well correlated (0.2586 < *r* < 0.3049) across the three strains and in particular for extreme values (**Supplemental Fig S1**), suggesting similar sequence content across strains and a minimal effect of experimental error. In addition, our results suggest that there are no genes found on the house fly X chromosome that are not present on the Y chromosome.

To ensure that our results are not an artifact of incomplete annotation of house fly X-linked genes, we calculated 
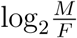
 coverage across non-overlapping 1 kb intervals in the reference genome. The distribution of 
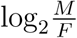
 across autosomes is expected to be centered at zero. If males have a single copy of the X chromosome, we should observe a second peak at 
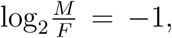
 indicating a 2-fold enrichment of X-linked sequences in females. We do indeed observe that the distributions of 
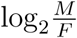
 are centered near zero for all three house fly strains in our analysis, but there is no obvious secondary peak at 
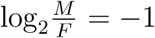
 in any of the distributions (Fig 2). To test for a secondary peak at

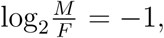
 we fit a mixture of two normal distributions to our data using an expectation-maximization algorithm with starting values of −1 and 0 for the means of the two distributions (Benaglia et al., 2009). Most of the 1 kb intervals (93–99%) are assigned to a distribution that is centered near zero, and the remainder of the intervals are assigned to a secondary distribution with a mean <0 (**Fig 2**). However, those secondary distributions all have a mean > −1, suggesting that there are few sequences present on the X chromosome at twice the abundance as on the Y. Additionally, 
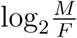
 for the 1 kb intervals is positively correlated (0.1661 < *r* < 0.2199) across the three strains, and in particular for extreme values (**Supplemental Fig S2**).

**Figure 2:**
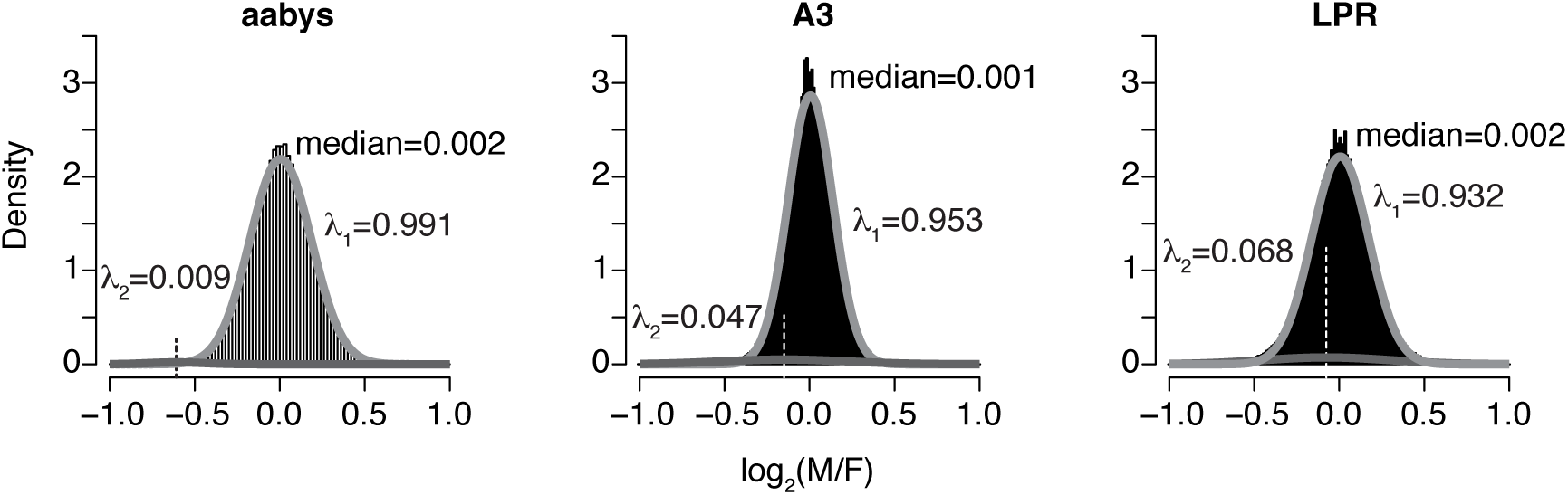
Histograms are plotted of 
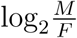
 for 1 kb intervals across three strains. The medians of the distributions are shown. Gray lines represent two normal distributions fit to the data from each strain. The λ_1_ values are the proportion of observed data estimated to be part of the normal distribution centered near zero, and λ_2_ is the proportion estimated to be part of the distribution with a mean <0. The dashed lines show the means of the distributions with λ_2_.

We next sought to identify Y-linked sequences that are absent from the X chromosome (i.e., the reciprocal of the analyses described above). To this end, we first used the male sequencing reads from the aabys strain to assemble a genome that contains a Y chromosome using SOAPdenovo2 (Luo et al., 2012). It was necessary to assemble a male genome because the genome project sequenced gDNA from female flies (Scott et al., 2014). Then we used a *k*-mer comparison approach to identify male-specific sequences by searching for male genomic scaffolds that are not matched by female sequencing reads (Carvalho and Clark, 2013). Most of the scaffolds in the male genome assembly were (nearly) completely matched by female sequencing reads, and none of the male scaffolds were completely unmatched by female sequencing reads (Fig 3). We obtain similar results when we use a male genome assembled with ABySS (Simpson et al., 2009) or if we assemble the male genome with SOAPdenovo2 using only male reads that do not align to the female assembly (**Supplemental Fig S3**). In contrast, when this approach was used to identify Y-linked scaffolds in species with differentiated sex chromosomes (*Drosophila* and humans), a substantial number of Y-linked scaffolds were completely unmatched by female sequencing reads (Carvalho and Clark, 2013). These results suggest that there are not large segments of the House fly Y chromosome that are unique from the X chromosome or the rest of the genome.

**Figure 3:**
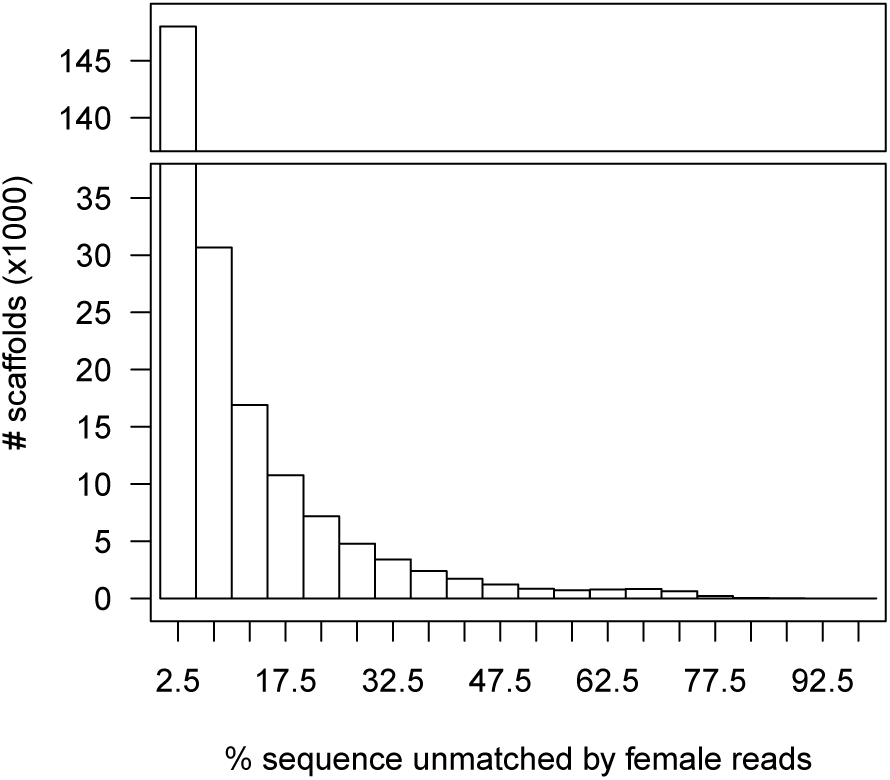
Histogram of female read mapping coverage to scaffolds assembled from a male genome.

We examined the scaffolds from the male house fly genome with a high percentage of sequence unmatched by female reads. We performed blastx searches of the 50 scaffolds with the highest percent unmatched sequence (79.7–96.6%) against the NCBI non-redundant protein database (Altschul et al., 1997). Only 6/50 scaffolds had hits to annotated house fly genes, whereas 27 had hits to transposable element (TE) sequences, 3 hit other sequences from other species, and 14 had no hits in the database (**Supplemental Data S4 & S5**). The scaffold with the highest percent unmatched sequence (96.6%) is 1,143 nucleotides long and contains a 219 basepair segment that matches an annotated house fly gene on chromosome 1 that is homologous to a *Drosophila melanogaster* gene with a predicted membrane associated GRAM domain (CG34392). This scaffold does not have any blastn or blastx hits to other sequences in the database. In addition, there are 22 scaffolds that are both >5 kb and >50% unmatched by female reads (**Supplemental Data S4 & S6**). We also performed blastx of those scaffolds against the NCBI database, and we found that 15/22 of hit a TE, 4 hit an annotated house fly gene, and 3 hit an other sequence from another species. None of the annotated house fly genes hit by these 72 scaffolds are predicted to be on element F (the house fly X chromosome). Notably, most of these scaffolds (42/72) have sequence similarity with a TE, suggesting that the Y chromosome may contain unique repetitive sequences or be enriched for particular repeat classes.

### 2.2. Moderate differences in sequence abundance between House fly males and females

We next examined whether housefly X and Y chromosomes exhibit differential representation of shared sequences, as might be expected from expansion or contraction of satellite repeats or other repetitive elements. We first used a principal components (PC) analysis to compare read mapping coverage of the male and female sequencing libraries across non-overlapping 1 kb intervals in the reference (XX female) genome. The first PC (PC1) explains 81.5–91.1% of the variance in coverage across libraries in the three strains, and PC1 clearly separates the male and female sequencing libraries in all three strains (**Fig 4**). Therefore, house fly males and females, and by association X and Y chromosomes, exhibit systematic differences in the abundance of some sequences.

**Figure 4:**
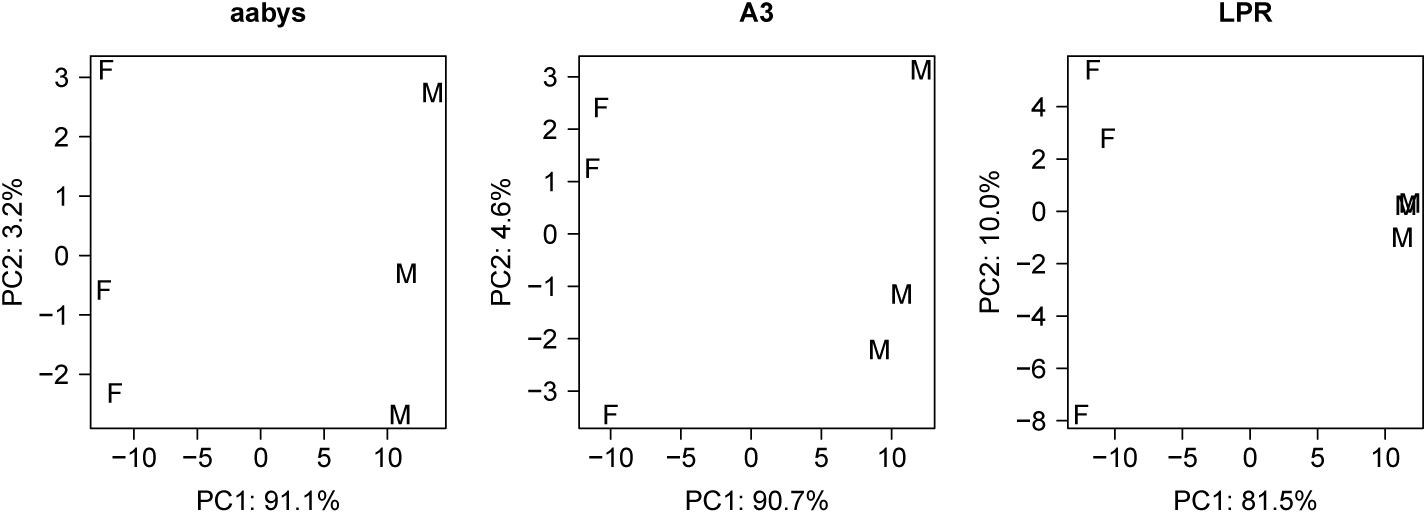
Plot of the first two principal components explaining differential sequencing coverage between female (F) and male (M) libraries.

We applied two different approaches to characterize sequences enriched on the X and Y chromosomes (i.e., differentially abundant in female and male genomes). First, we searched for 1 kb windows with significantly different coverage between males and females (false discovery rate corrected *P* < 0.05 and 
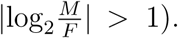
 We identified 214 of these “sex-biased” windows: 63 are >2-fold enriched in females, and 151 are >2-fold enriched in males (**Supplemental Data S7**). The X and Y chromosomes of house fly are largely heterochromatic (Boyes et al., 1964; Hediger et al., 1998b), and it is possible that differences in the abundances of particular repetitive DNA sequences (e.g., TEs and other interspersed repeats) between the X and Y chromosomes are responsible for the differences in read coverage between females and males. Sequences from repetitive heterochromatic regions of the genome are less likely to be mapped to a genomic location (Smith et al., 2007), and we therefore expect sex-biased windows to be located on scaffolds that are not mapped to a house fly chromosome. Only 2/63 (3.2%) female-enriched windows are within a scaffold that we were able to map to a chromosome (neither was mapped to element F, the ancestral X chromosome). In addition, 59/151 (39.1%) male-enriched windows are within a scaffold that maps to a Muller element (only one of those scaffolds maps to element F). In contrast, 65.7% of 1 kb windows that are not differentially covered between males and females are on scaffolds that we are able to map to Muller elements (2033/3096 windows with *P* > 0.05 and 
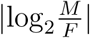
 < 0.01). These unbiased windows are more likely to be mapped to a Muller element than the sex-biased windows (*P* < 10^−15^ in Fisher’s exact test), providing some evidence that differential coverage between males and females could be driven by repeat content differences between the X and Y chromosomes.

We next tested for an enrichment of annotated repeats within the female- and male-biased 1 kb windows, and we found that all 63 of the female-biased windows and most of the male-biased windows (149/151) contain sequences masked as repetitive during the house fly genome annotation (Supplemental Data). However, 3071/3096 (>99%) of the 1 kb windows that are not differentially covered between males and females also contain repeat masked sequences; this fraction is not significantly different than the fraction of repeat masked sex-biased windows (*P* = 1 for female-biased and *P* = 0.6 for male-biased windows using Fisher’s exact test). In addition, the proportion of sites within male-biased and female-biased windows that are repeat masked is less than that of unbiased windows, suggesting that the sex-biased windows are actually depauperate for annotated repeats (**Supplemental Fig S4**). However, these analyses are limited because a large fraction (≥ 52%) of the house fly genome is composed of interspersed repeats that are poorly annotated (Scott et al., 2014). Future improvements to repeat annotation in the housefly genome may therefore shed light on the nature of repetitive sequences that differentiate the X and Y chromosomes.

**Figure 5:**
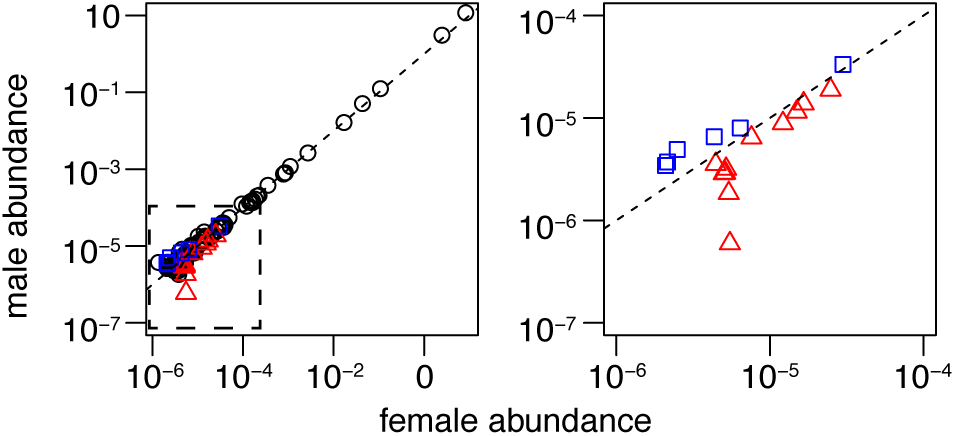
Minimal differences between *k*-mer abundances in male and female genomes. The left plot shows the abundances of the 100 most common *k*-mers in the male and female sequencing reads averaged across the three libraries from each sex in the aabys strain. The dashed box in the left graph indicates the subset of the range plotted on the right graph, which contains only *k*-mers where abundances in all three libraries from one sex are greater than the three libraries from the other sex. Triangles indicate *k*-mers where abundances in all three female libraries are greater than the three male libraries, and squares indicate k-mers that are more abundant in male libraries. The dashed line indicates equal representation in males and females.

As a second approach to identify candidate X- or Y-enriched sequences, we first determined the abundances of all possible 2–10mers in the male and female aabys sequencing reads. This approach will identify smaller sequence motifs that may differentiate the X and Y chromosomes than the analysis described above, and it does not require any *a priori* repeat annotations. The 100 most common *k*-mers are found at similar frequencies in both males and females (Fig 5), with the abundances highly correlated between sexes (*r* = 0.999). We considered a *k*-mer to be over-represented in one sex if the minimum abundance across the three replicate libraries for that sex is greater than the maximum in the other sex. Six *k*-mers are over-represented in males using this cutoff, but they are all less than 2-fold enriched in males (**Fig 5 & Supplemental Fig S5**). These results suggest that short sequence repeats do not predominantly differentiate the X and Y chromosomes.

### 2.3. Relative heterozygosity in males and female suggests that the house fly Y chromosome is very young

Our data suggest that, other than the unidentified M factor, the house fly X and Y chromosomes do not contain any genes not found on the gametologous sex chromosome, and we find little evidence for unique sequences on the X or Y. We therefore hypothesize that the house fly Y chromosome is actually an ancestral brachyceran X chromosome that recently acquired an M factor. While recently derived neo-Y chromosomes may not differ in gene content from the gametologous X chromosome, modest sequence-level X-Y differentiation can result in elevated heterozygosity within sex chromosome genes in males (Vicoso and Bachtrog, 2015). We tested for elevated heterozygosity by first identifying polymorphic sites (SNPs) within genes in aabys males and females. Heterozygosity is elevated in X chromosome (element F) genes relative to autosomes in both males and females (**Supplemental Fig S6**). However, when we compare the proportion of heterozygous SNPs in males relative to females for genes on each chromosome (**Fig 6A**), genes on the X chromosome resemble autosomal genes with equivalent heterozygosity in males and females (*P* = 0.45 in a Mann-Whitney test comparing male: female heterozygosity on element F with the other chromosomes). This result demonstrates that the house fly Y is so young that Y chromosome genes have not yet accumulated modest sequence differences from the X chromosome.

Some house fly males carry the M factor on the third chromosome (III^M^), and they have two copies of the X chromosome without an M factor (Hamm et al., 2015). The III^M^ chromosome is a recently derived neo-Y not found in close relatives, and we expect that males heterozygous for III^M^ (hereafter III^M^ males) will have an excess of heterozygous SNPs on the third chromosome. To test this hypothesis, we used available RNA-Seq data (Meisel et al., 2015) to calculate the proportion of heterozygous SNPs in III^M^ males relative to XY males (Fig 6B). As predicted, there is an excess of heterozygous SNPs on the third chromosome in III^M^ males relative to XY males (*P* = 10^−122^) in a Mann-Whitney test comparing chromosome III with the other autosomes). III^M^ males also have an elevated number of strain-specific SNPs on the third chromosome (**Supplemental Fig S7**). Surprisingly, there is also increased heterozygosity on the X chromosome in III^M^ males relative to XY males (*P* = 10^−4^) even though III^M^ males have the XX genotype. We also observe elevated heterozygosity on the third chromosome and X chromosome in III^M^ males relative to females of the same strain, but not on the X chromosome in XY males relative to XX females (Supplemental Fig S8). These results further support our conclusion that house fly Y chromosome genes are not differentiated from X chromosome genes. In contrast, the III^M^ chromosome harbors evidence that it is partially differentiated from the non-M-bearing third chromosome.

**Figure 6:**
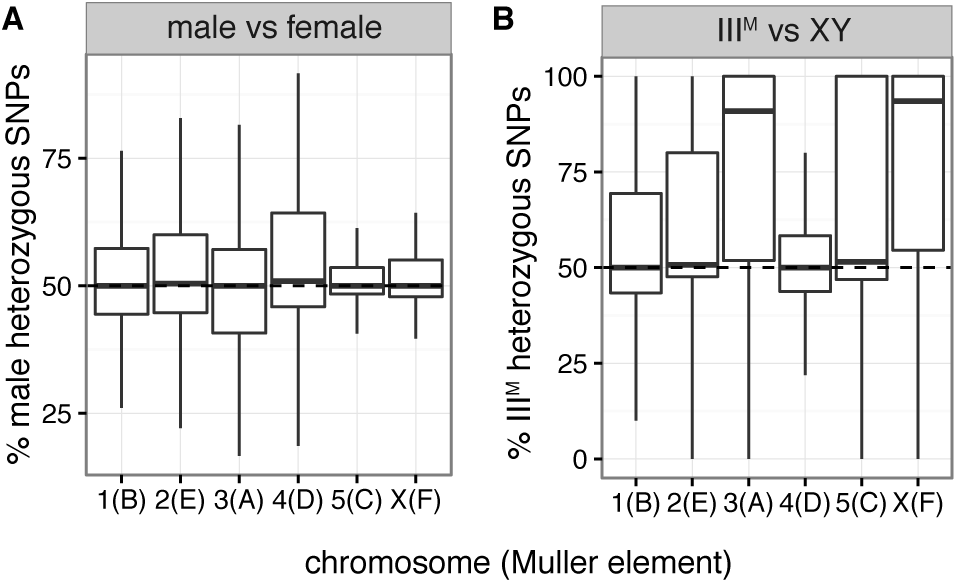
There is elevated heterozygosity on the third chromosome in III^M^ males, but not on the X chromosome in XY males. Box plots show the distribution of the percent of heterozygous SNPs within genes on each chromosome in either (A) XY aabys males relative to XX aabys females using genomic DNA sequences or (B) III^M^ males relative to XY males using RNA-seq data. Values >50% indicate elevated heterozygosity in XY males or III^M^ males. The median across all autosomes is indicated by a dashed line.

## 3. Discussion

We found very little evidence for differentiation between the X and Y chromosomes in house fly, despite the fact that House fly has a karyotype that resembles the ~100 my old ancestral brachyceran karyotype (Boyes et al., 1964; Foster et al., 1981; Weller and Foster, 1993; Vicoso and Bachtrog, 2013). There are few sequences unique to the X or Y (Figs 1, 2, & **3**), little evidence for differential abundance of specific sequences on the X and Y (Fig 5, but see Fig 4), and no elevated heterozygosity within X chromosome genes in males (Fig 6). We conclude that the house fly X and Y chromosomes do not contain different genes (other than M), which is consistent with previous experiments that failed to identify sex-linked markers and found that XX, XO, and YO flies are viable and fertile (Bull, 1983; Hediger et al., 1998a; Hamm et al., 2015). Additionally, XY males have equal or greater expression of X chromosome genes when compared to XX (III^M^) males (Meisel et al., 2015), providing further evidence that XY males do not have a haploid X chromosome dose. In summary, our results suggest that the house fly Y chromosome is an ancestral brachyceran X chromosome that very recently acquired an M factor.

### 3.1 X-Y dierentiation in house fly

We failed to identify the male-determining M factor on the house fly Y chromosome. Previous *in situ* hybridizations of chromosomal dissections to mitotic chromosomes detected Y-specific, but not X-specific, segments of the house fly genome (Hediger et al., 1998b). The sequences of these chromosomal segments are unknown, but they presumably include the yet to be identi ed M factor. We hypothesize that our failure to identify M and other Y-specific regions is because they are small relative to the rest of the chromosome and/or di cult to assemble using short sequencing reads.

Despite the lack of genic differentiation between the house fly X and Y chromosomes (other than M), there are morphological differences between the X and Y that can be identifed through cytological examination (Boyes et al., 1964). Our results suggest that some of the morphological differences between the X and Y chromosomes result from the differential abundance of particular sequences between X and Y rather than the extensive differentiation that characterizes ancient pairs of sex chromosomes (Fig 4). Similarly, cytological analyses previously characterized separate X chromosomes carrying M (X^M^) that were thought to be different from Y^M^ (Denholm et al., 1983; Cakir and Kence, 1996). Our results suggest that Y^M^ and X^M^ chromosomes are morphological variants of the same neo-Y chromosome that arose when an ancestral X chromosome acquired an M factor. Such morphological or repeat content variation in Y chromosomes has been previously documented in *Drosophila* and other dipterans (Dobzhansky, 1935; Miller and Stone, 1962; Miller and Roy, 1964; Lyckegaard and Clark, 1989; Lemos et al., 2008; Hall et al., 2016).

### 3.2. Creation of house fly neo-Y chromosomes

We hypothesize that an M factor recently translocated to the house fly X chromosome to create a neo-Y, and the ancestral Y chromosome was lost from house fly populations (Fig 7). Alternatively, the house fly Y chromosome could have arisen through the fusion of the ancestral Y and X chromosomes. However, after an X-Y fusion, the neo-Y should retain ancestral Y-specific sequences, which we fail to detect. The III^M^ chromosome is also a neo-Y that likely arose when an M factor translocated onto a standard third chromosome (Fig 7). Curiously, III^M^ males have elevated heterozygosity in X chromosome genes relative to XX females and XY males (**Figs 6 & S8**). One possible cause of this elevated heterozygosity is that some X chromosome genes were translocated onto III^M^ along with the M factor. The elevated X chromosome heterozygosity we detect in III^M^ males would therefore be the result of those males being triploid for X chromosome genes. No nullo-X/Y flies carrying III^M^ have been identi ed to our knowledge, suggesting that the X/Y chromosome contains some essential genes not translocated to III^M^. Additional work is necessary to determine the nature of the translocations that created the Y and III^M^ chromosomes.

**Figure 7:**
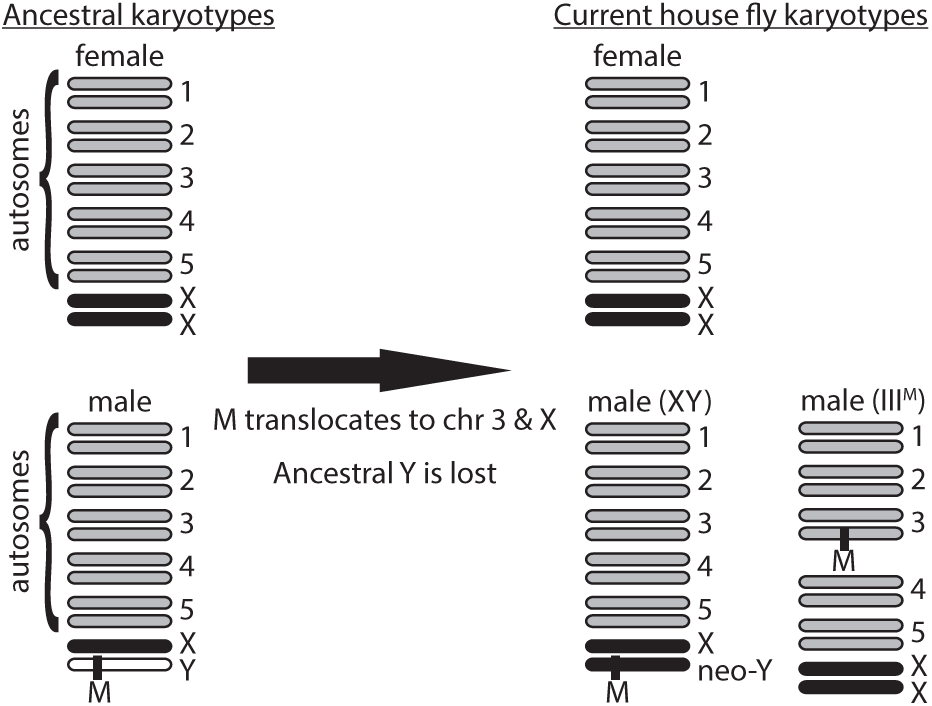
Ancestral and hypothesized derived house fly karyotypes.

Our results are also suggestive of the order of events that created the Y and III^M^ chromosomes. III^M^ males have elevated heterozygosity on the third chromosome, whereas XY males do not have elevated heterozygosity on element F (Fig 6). This suggests that the Y chromosome is a younger neo-Y than III^M^, but there is an alternative explanation for the patterns of polymorphism. differentiation between nascent X and Y chromosomes is accelerated by suppressed recombination in XY males (Charlesworth, 1991; Rice, 1996; Charlesworth et al., 2005; Bachtrog, 2013). Lack of recombination can be an inherent property of male meiosis (as in *Drosophila*) or arise via Y chromosome inversions that suppress crossing over between the X and Y. There is evidence for male recombination in house flies (Feldmeyer et al., 2010), suggesting that chromosomal inversions would be required for recombination suppression between the X and Y. If the III^M^ chromosome carries inversions and the Y chromosome does not, then the elevated heterozygosity in III^M^ males but not XY males could be a result of inversions accelerating the rate of divergence between the III^M^ chromosome and the standard third chromosome (Navarro et al., 2000). Additional sequencing of XY and III^M^ males is needed to test for inversions.

### 3.3. Translocating sex-determining loci can cause sex chromosome recycling and create cryptic neo-sex chromosomes

Our results provide the first evidence, to our knowledge, of the conversion of an existing X chromosome into a Y chromosome (or Z into W), recycling a differentiated sex chromosome pair into nascent sex chromosomes without any evidence of fusion to an autosome. In comparison, most previously documented sex chromosome transitions involved autosomes transforming into sex chromosomes through either the evolution of a novel sex determining locus on the autosome or a fusion of the autosome with a sex chromosome (e.g., Patterson and Stone, 1952; Steinemann and Steinemann, 1998; Filatov et al., 2000; Liu et al., 2004; Veyrunes et al., 2004; Carvalho and Clark, 2005; Vallender and Lahn, 2006; Ross et al., 2009; Vicoso and Bachtrog, 2013; Bachtrog et al., 2014; Beukeboom and Perrin, 2014; Vicoso and Bachtrog, 2015). There are other examples of sex chromosome transformations involving only X, Y, Z, and W chromosomes (i.e., no autosomes) in platyfish, *Rana rugosa*, and *Xenopus tropicalis* (Kallman, 1984; Miura, 2007; Roco et al., 2015). These X/Y/Z/W transformations in fish and frogs, however, involve nascent sex chromosomes, not ancient sex chromosomes as in house fly. Moreover, the sex chromosome transitions in platyfish, *R. rugosa*, X. and *tropicalis* all involve a change in the heterogametic sex (i.e., XY males to ZW females, or vice versa), whereas the house fly X and Y chromosomes did not switch to a Z and W.

We hypothesize that the X-to-Y conversion in house fly occurred because the male-determining locus translocated from the Y to the X. Translocating sex determining loci are rare and do not typically include long-established sex chromosomes (Traut and Willhoeft, 1990; Woram et al., 2003; Faber-Hammond et al., 2015), suggesting that X-to-Y (or Z-to-W) conversion similar to house fly may not be observed in other taxa. However, there is rampant gene traffic to and from long-established Y chromosomes (Koerich et al., 2008; Hughes et al., 2015), providing a possible mechanism for the Y-to-X (or W-to-Z) translocation of a sex determining locus in other taxa even if the sex determiner does not exhibit a high rate of translocation on its own. The fact that the neo-Y chromosome in house fly remained undetected despite decades of work on this system (Dübendorfer et al., 2002) suggests that X-to-Y transitions may have occurred in other taxa and remain cryptic because the karyotype has remained unchanged.

## 4. Methods

### 4.1. Fly strains

We used five house fly strains to identify X-and Y-linked sequences. One strain, Cornell susceptible (CS), has been reported to have X/X; III^M^/III males (Scott et al., 1996; Hamm et al., 2005; Meisel et al., 2015). The other four strains have previously been characterized as having males with the XY karyotype: aabys, A3, LPR, and CSaY. The genome strain, aabys, has recessive phenotypic markers on each of the five autosomes (chromosomes I–V and had been cytologically determined to have XY males (Wagoner, 1967; Tomita and Wada, 1989; Scott et al., 2014). The A3 strain was generated by crossing XY males from a pyrethroid-resistant strain (ALHF) with aabys females (Liu and Yue, 2001). The LPR strain is a pyrethroid-resistant strain that was previously determined to have XY males (Scott and Georghiou, 1985; Scott et al., 1996). Finally, the CSaY strain was created by crossing aabys males (XY) with CS females, and then backcrossing the male progeny to CS females to create a strain with the aabys Y chromosome on the CS background (Meisel et al., 2015). We validated that the M factor is not on an autosome in the A3, LPR, and CSaY strains by crossing males of each strain to aabys females, and then we backcrossed the male progeny to aabys females. We did not observe sex-linked inheritance of any of the aabys phenotypic markers, confirming that the M factor is not on chromosomes I–V in A3, LPR, or CSaY. Females of all strains were expected to be XX.

### 4.2. Genome sequencing, mapping, and assembly

The house fly genome consortium sequenced, assembled, and annotated the genome using DNA from female flies of the aabys strain, a line with XX females and XY males (Scott et al., 2014). The annotation includes both predicted genes and inferred homology relationships with *D. melanogaster* genes, and we used the orthology calls from annotation release 100 (version 2.0.2) to assign house fly genomic scaffolds to chromosome arms using a majority rule as described previously (Meisel et al., 2015). Briefly, scaffolds were assigned to a Muller element if the majority of genes on the scaffold with 1:1 *D. melanogaster* orthologs have orthologs on the same D. melanogaster element. In total, 62 house fly genes have 1:1 *D. melanogaster* orthologs on Muller element F, which amounts to 3/4 of the ~80 genes on *Drosophila* element F (Leung et al., 2010). We used these 1:1 orthologs to assign seven house fly scaffolds to element F (the X chromosome), and those seven scaffolds contain 51 genes. We repeated all of the analyses described in the Results using only genes with 1:1 *D. melanogaster* orthologs and obtained qualitatively similar results as when we used scaffold-level Muller element assignments.

We sequenced genomic DNA (gDNA) from aabys male and female heads with 150 bp paired-end reads on an Illumina NextSeq 500 at the University of Houston genome sequencing core. Three replicate libraries of each sex were prepared using the Illumina TruSeq DNA PCR-free kit, and the six libraries were pooled and sequenced in a single run of the machine. We also sequenced gDNA from three replicates of male and female heads from A3 and LPR flies (12 samples total) in a single run on the NextSeq 500 using 75 bp paired-end reads. For each of the 18 sequencing libraries, DNA was extracted from separate pools of fly heads using the QIAGEN DNeasy Blood & Tissue Kit. Illumina sequencing reads were mapped to the assembled house fly genome using BWA-MEM with the default parameters (Li and Durbin, 2009; Li, 2013), and we only included uniquely mapping reads where both ends of a sequenced fragment mapped to the same scaffold in the reference genome. Reads that failed to meet these criteria were considered unmapped for the male genome assembly described below. Mapping statistics are presented in **Supplemental Table S1**.

We additionally assembled the reads from aabys male samples using SOAPdenovo2 (Luo et al., 2012) and ABySS (Simpson et al., 2009) to construct a reference genome that contains Y-linked sequences. Mapping our sequence data to the reference genome revealed that our average insert size was 370 bp (**Supplemental Fig S9**), which was used as a parameter in the SOAPdenovo2 genome assembly, along with a pair number cutoff 3 and a minimum alignment length 32 bp. For the ABySS assembly we used a *k*-mer pair span (k) of 64. We also assembled a genome from only male reads that did not align to the female genome reference assembly using SOAPdenovo2 (Luo et al., 2012). For downstream analyses, we only retained scaffolds with a length ≥1000 bp in each assembly. Assembly statistics are presented in **Supplemental Table S2**.

### 4.3. Identifying X-and Y-linked sequences

We used four differential coverage approaches to identify candidate X-and Y-linked sequences in the house fly genome. The first approach identifies X-linked genes or sequences by testing for 2-fold higher abundance in females relative to males (Vicoso and Bachtrog, 2013). To do this, we used DESeq2 to calculate the log_2_ relative coverage within individual genes and 1 kb windows between the three male and female derived libraries (Love et al., 2014). We also used DESeq2 to calculate P-values for differential coverage between females and males.

The second approach was used to identify Y-linked sequences by searching for scaffolds in the male genome assembly that are missing from the female sequencing reads. We only considered assembled scaffolds from the male genome that were ≥1 kb. We implemented a *k*-mer comparison approach to identify male-specific sequences (Carvalho and Clark, 2013). In our implementation, we used a *k*-mer size of 15 bp, used the male sequencing reads to construct a validating bit-array, and implemented the options described by Carvalho and Clark (2013) for identifying Y-linked sequences in *Drosophila* genomes (**Supplemental Methods S1 & S2**).

In the third approach, we analyzed gDNA sequencing reads from aabys males and females to identify *k*-mers with sexually dimorphic abundances. We used the k-Seek method to count the abundance of 2-10mers in the three male and three female aabys sequencing libraries (Wei et al., 2014). We normalized the *k*-mer counts by multiplying the count by the length of the *k*-mer and dividing by the number of reads in the library.

The fourth approach identifies nascent sex chromosomes because they have elevated heterozygosity in the heterogametic sex (Vicoso and Bachtrog, 2015). We implemented this approach using both gDNA-and mRNA-Seq data. For the gDNA-Seq, we used the Genome Analysis Toolkit (GATK), following the best practices provided by the software developers (McKenna et al., 2010). Starting with the male and female mapped reads from the aabys strain described above, we identified duplicate reads. Insertions and deletions (indels) were identified and realigned using RealignerTargetCreator and IndelRealigner, respectively. We then called variants in each of the six aabys sequencing libraries using HaplotypeCaller, and we selected the highest quality SNPs and indels using SelectVariants and VariantFiltration (for SNPs: QD < 2, MQ < 40, FS > 60, SOR > 4, MQRankSum < −12.5, ReadPosRankSum < −8; for indels: QD < 2, ReadPosRankSum < −20, FS > 200, SOR > 10). The high quality SNPs and indels were next used for recalibration of the base calls with BaseRecalibrator and PrintReads. The process of variant calling and base recalibration was performed three times, at which point there were no benefits of additional base recalibration as validated with AnalyzeCovariates. We next used the recalibrated reads from all three replicates of each sex to call variants in males and females using HaplotypeCaller with emission and calling confidence thresholds of 20. We filtered those variants using VariantFiltration with a cluster window size of 35 bp, cluster size of 3 SNPs, FS > 20, and QD < 2. We used the variant calls to identify heterozygous SNPs within genes using the coordinates from the genome sequencing project (Scott et al., 2014). An example script with our SNP calling pipeline is available in **Supplemental Methods S3.**

When we implemented the GATK pipeline for variant calling of the mRNA-Seq data (accession: GSE67065; Meisel et al., 2015), we used STAR to align reads from 6 XY male libraries and 6 III^M^ male libraries separately (Dobin et al., 2013). After aligning reads to the reference genome, we used the aligned reads to create a new reference genome index from the inferred spliced junctions in the first alignment, and then we performed a second alignment with the new reference. We next marked duplicate reads and used SplitNCigarReads to reassign mapping qualities to 60 with the ReassignOneMappingQuality read filter for alignments with a mapping quality of 255. Indels were realigned and three rounds of variant calling and base recalibration were performed as described above for the gDNA-Seq data. We applied GenotypeGVCFs to the variant calls from the 2 strains for joint genotyping of all samples, and then we used the same filtering parameters as used in the gDNA-Seq to extract high quality SNPs and indels from our variant calls.

## 5. Data Access

All sequence data have been submitted to GenBank under accession PRJNA342472.

## 6. Acknowledgements

This project was initiated during discussions with Andy Clark and Rob Unckless, who provided valuable comments throughout the completion of this work. Jeff Scott kindly supplied the A3 and LPR flies. Illumina sequencing was performed by the University of Houston Sequencing Core, with the assistance of Yinghong Pan and Utpal Pandya. Computational analyses were performed at the University of Houston Center for Advanced Computing and Data Systems, with some assistance from Adrian Garcia and Shuo Zhang. We thank Erin Kelleher for feedback on the preparation of this manuscript. This work was supported by start-up funds from the University of Houston.

## Supplemental Figures and Tables

**Figure S1:**
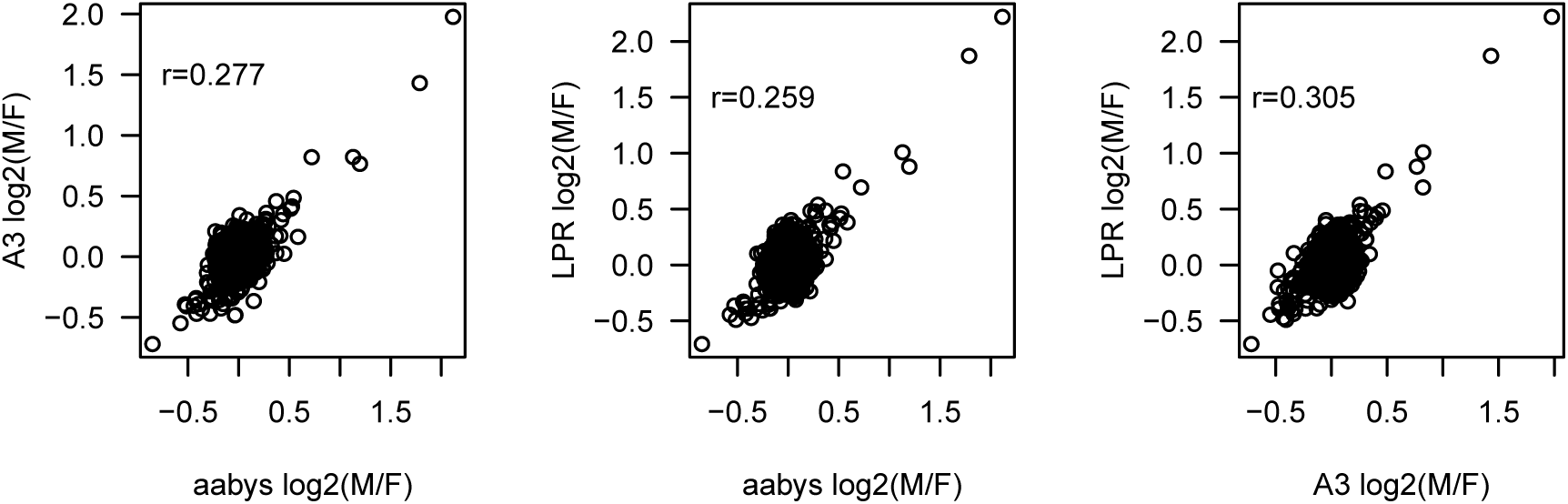
Correlation of 
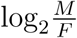
 for genes between three strains of House fly.

**Figure S2:**
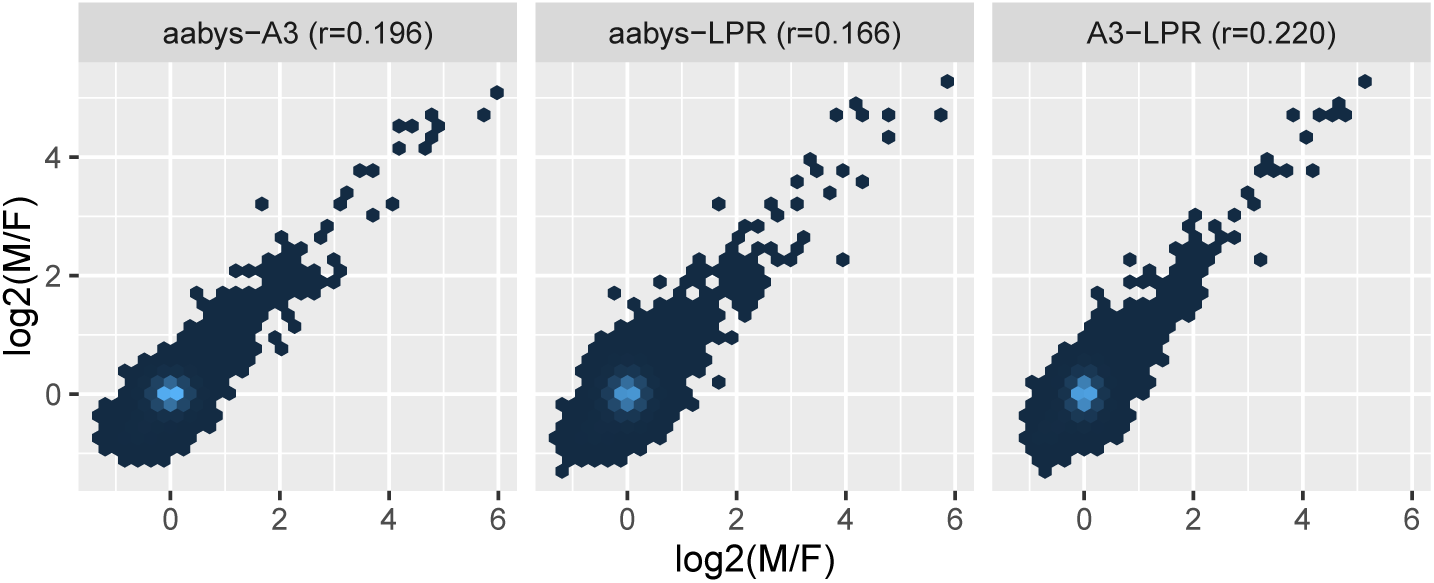
Correlation of 
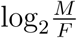
 for 1 kb intervals between three strains of house fly. 1 kb intervals are binned based on their 
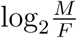
 values, with lighter colors indicating more intervals within a bin.

**Figure S3:**
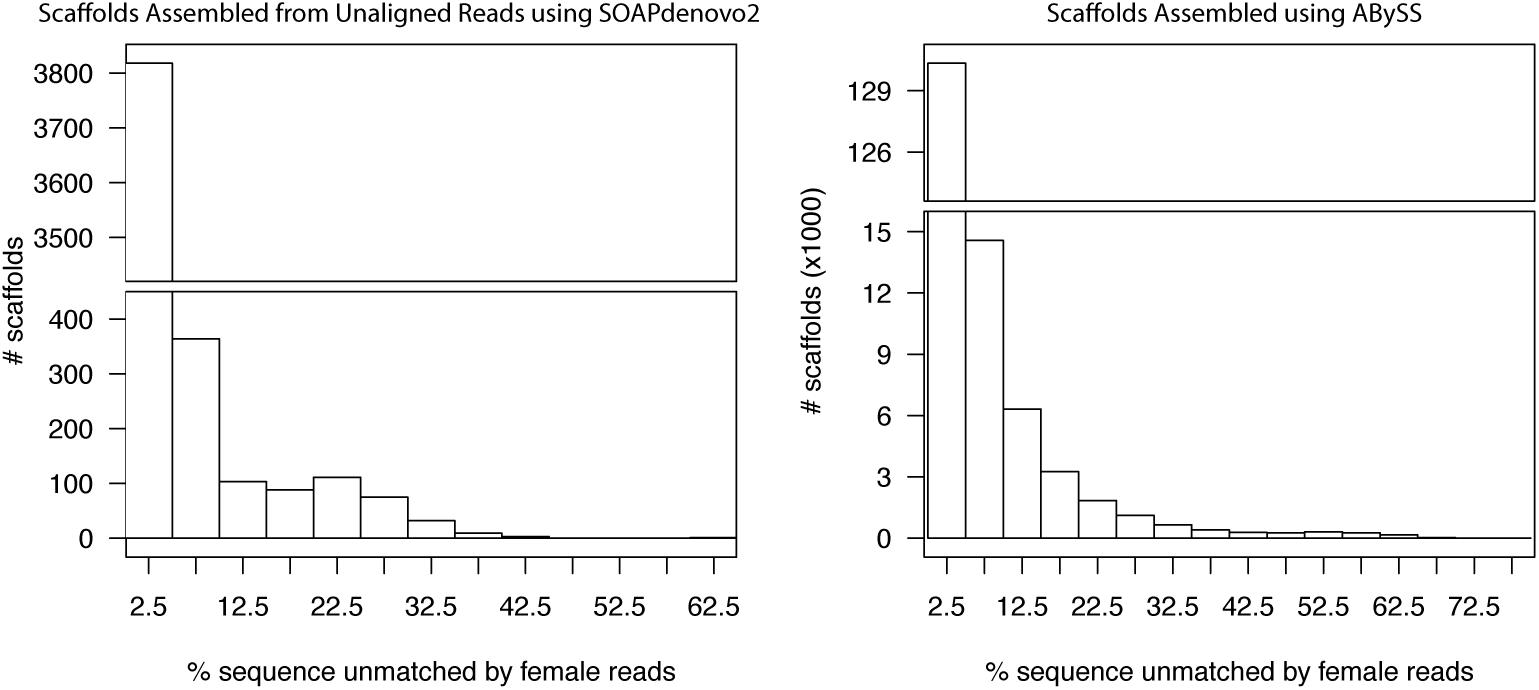
Histograms of female read mapping coverage to scaffolds from male genomes assembled using SOAPdenovo2 with reads that did not align to the female reference genome (left) or using ABySS (right).

**Figure S4:**
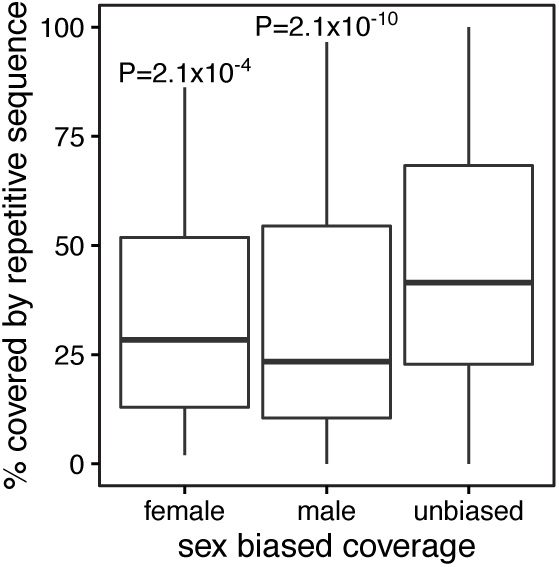
Boxplots show the distribution of the percent of 1 kb windows that contain predicted repetitive sequence. Three different types of 1 kb windows are plotted: those with female-biased read mapping coverage 
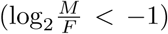
, those with male-biased coverage 
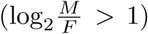
, and those with insignificant differences in coverage (unbiased). *P* values comparing the female- and male-biased windows with the unbiased windows from a Mann-Whitney test are shown.

**Figure S5:**
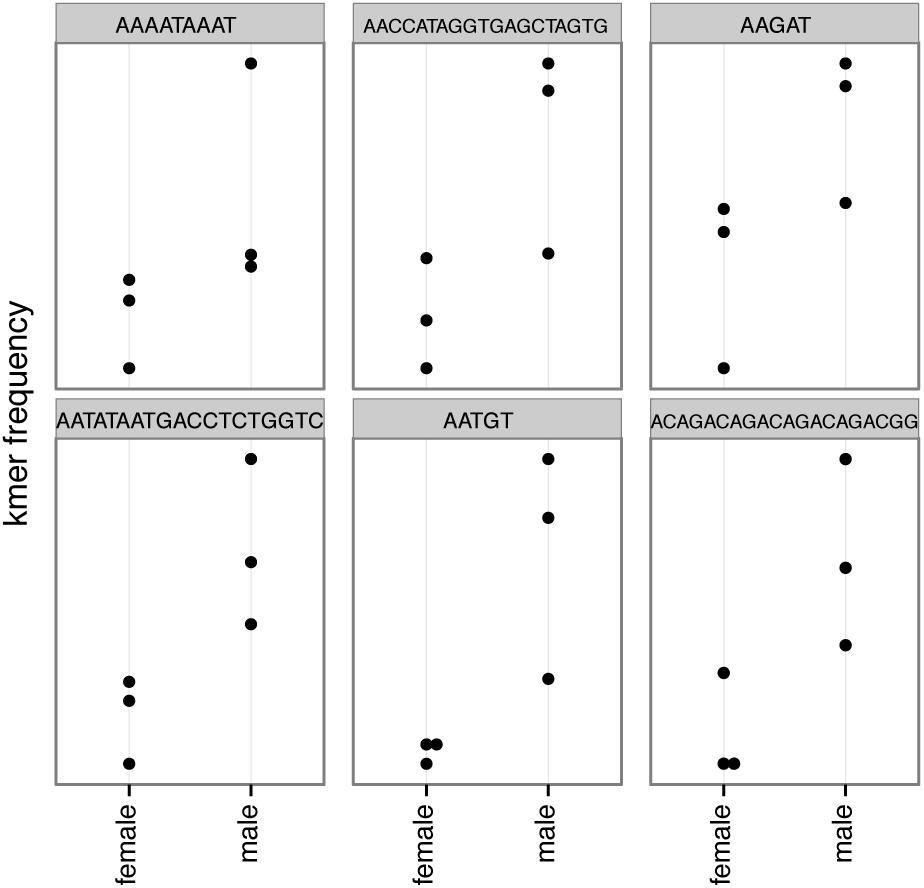
The frequency of the six *k*-mers over-represented in males is plotted for each of the 3 female and 3 male libraries.

**Figure S6:**
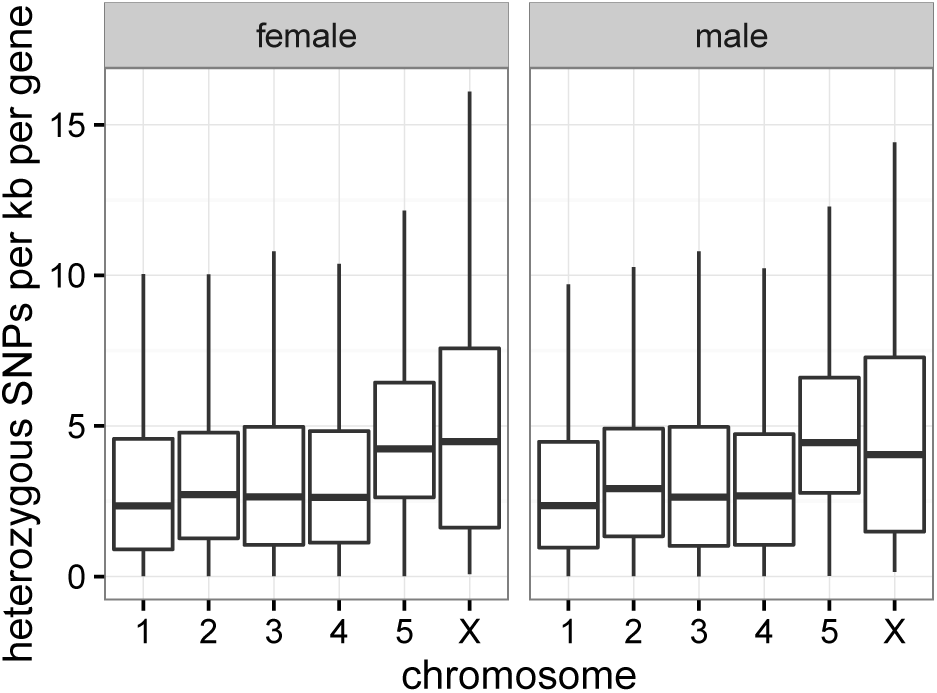
Boxplots show the distributions of heterozygous SNPs per kb per gene for each chromosome from the re-sequencing of aabys females and males.

**Figure S7:**
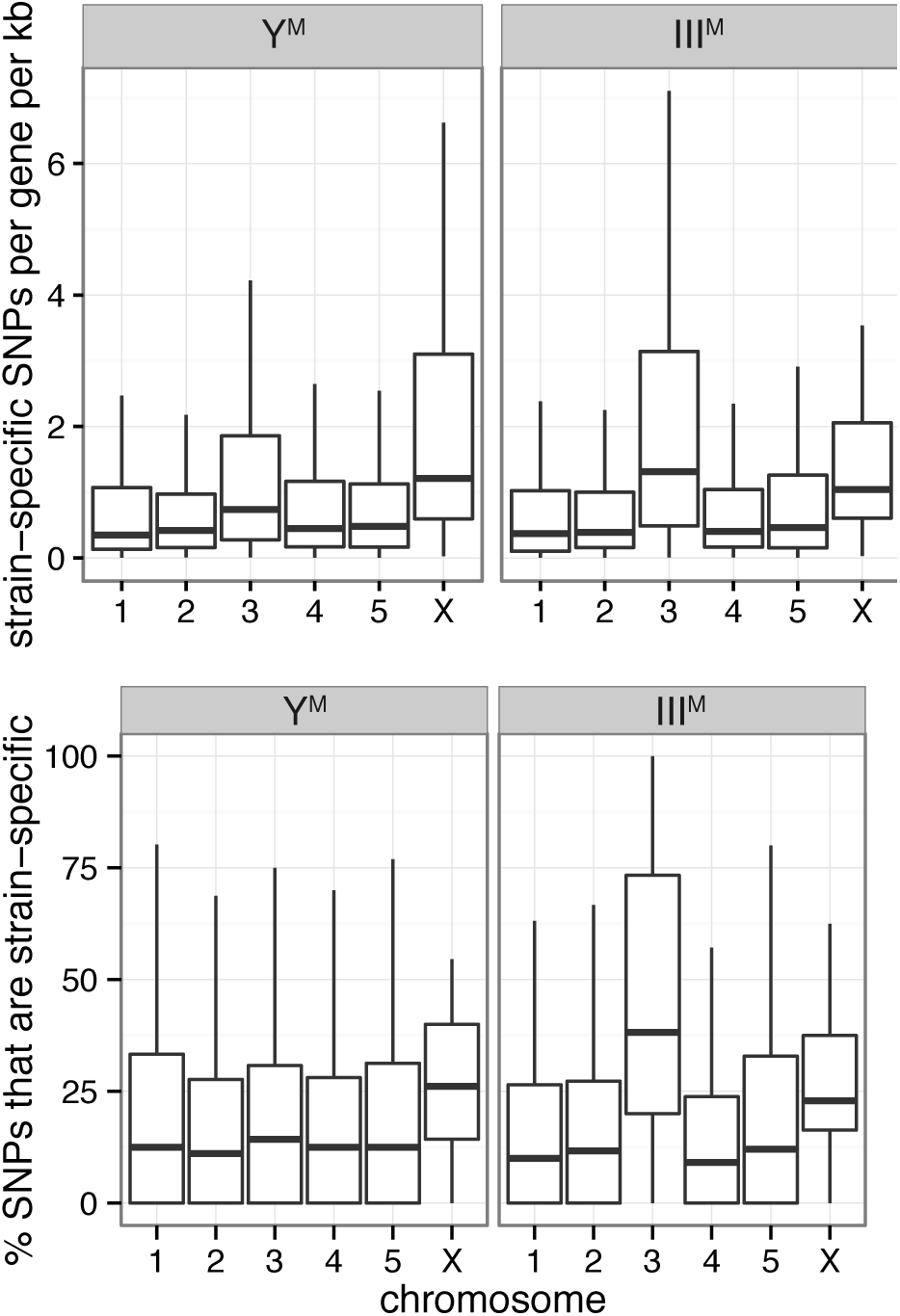
Boxplots show the distributions of strain-specific SNPs per gene per kb (top) and the percent of SNPs that are strain-specific per gene (bottom) for each chromosome from RNA-Seq data collected in Y^M^ and III^M^ males.

**Figure S8:**
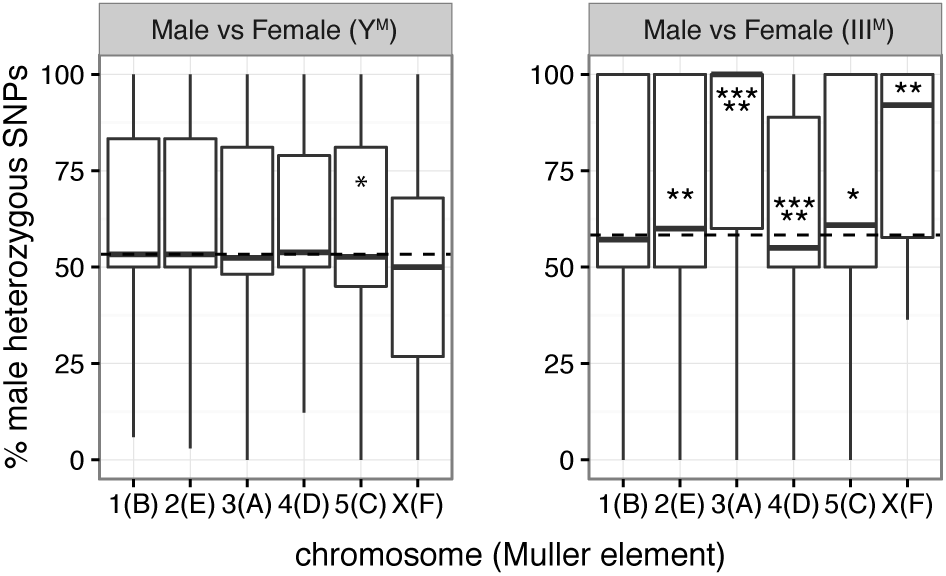
Box plots show the distributions of the percent of heterozygous SNPs within genes on each chromosome in males relative to females from strains with XY males (left) or III^M^ males (right). The median across all autosomes is indicated by a dashed line. Asterisks indicate significant differences between a chromosome and all other autosomes in a Mann-Whitney test (\**P* < 0.05, \*\**P* < 0.005, \*\*\*\*\**P* < 0.000005).

**Figure S9:**
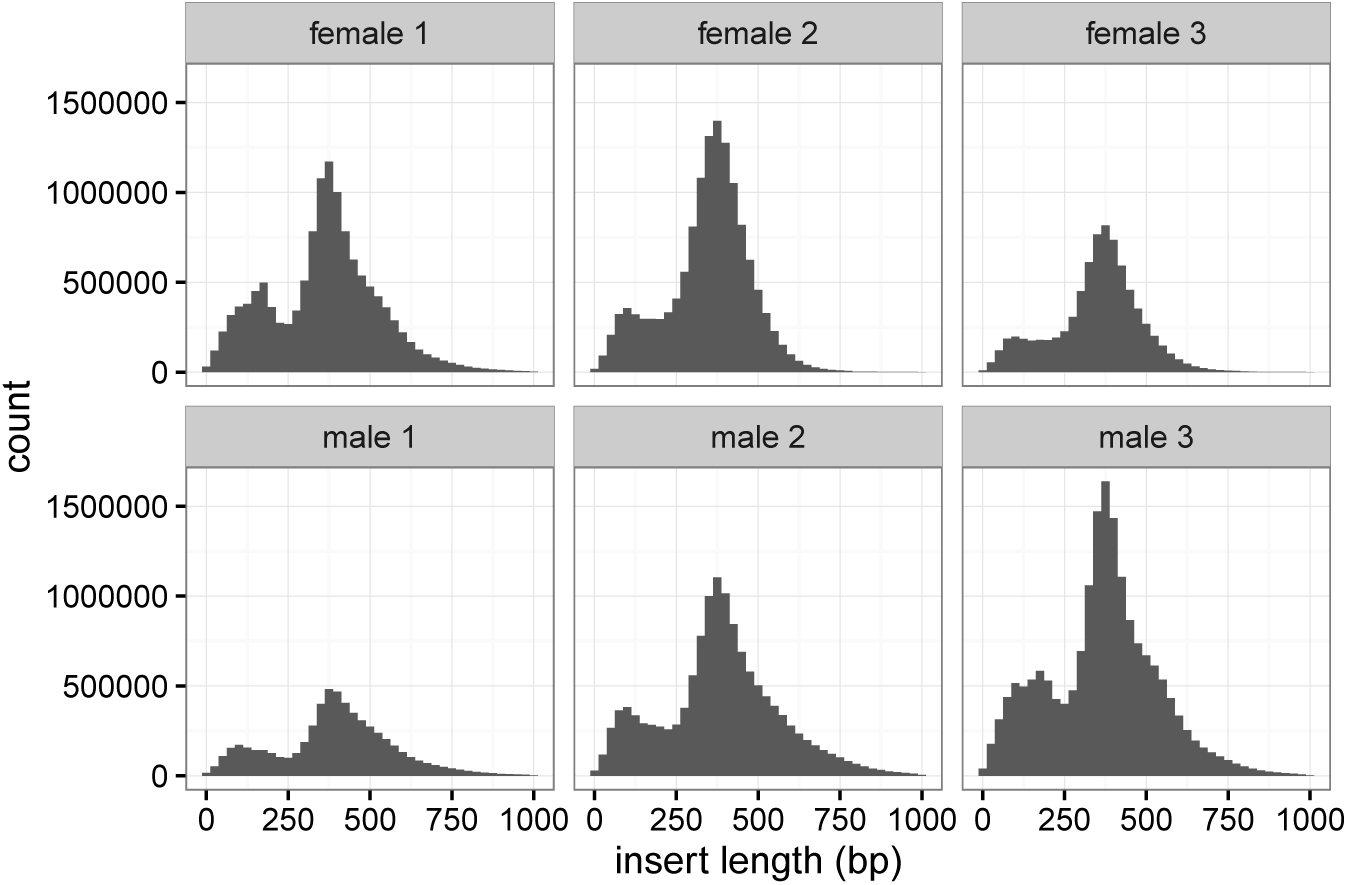
Distributions of insert sizes for the 6 male and female aabys libraries.

**Table S1.** Read mapping statistics for aabys, A3, and LPR male and female sequencing libraries. The number of paired reads that map to the same scaffold (mapped), where only one of two reads in a pair mapped (mapped_single), where two reads map to different scaffolds (mapped_distant), and where both reads failed to map (unmapped) are shown.

| strain | library | mapped | mapped_single | mapped_distant | unmapped |
| --- | --- | --- | --- | --- | --- |
| aabys | female1-26111291 | 13049678 | 887676 | 6292401 | 501177 |
| aabys | female2-26123107 | 13712814 | 809792 | 7125338 | 427100 |
| aabys | female3-26124109 | 7972753 | 550477 | 4716107 | 250508 |
| aabys | male1-26127107 | 6081087 | 656412 | 4333290 | 221249 |
| aabys | male2-26119223 | 13559671 | 1177151 | 8930649 | 514483 |
| aabys | male3-26113283 | 18323724 | 1275507 | 9500131 | 563536 |
| A3 | 1_A3_female_S10 | 21641219 | 2157816 | 5795725 | 1116445 |
| A3 | 2_A3_female_S7 | 22594865 | 2070878 | 6009122 | 1099204 |
| A3 | 3_A3_female_S6 | 28633153 | 2966799 | 8121340 | 1412310 |
| A3 | 4_A3_male_S9 | 22766977 | 2407270 | 6228309 | 1206439 |
| A3 | 5_A3_male_S5 | 21817355 | 2136473 | 5675312 | 1140114 |
| A3 | 6_A3_male_S4 | 19437196 | 1902804 | 4873879 | 1077971 |
| LPR | 7_LPR_female_S3 | 16238580 | 2325002 | 5586155 | 1062667 |
| LPR | 8_LPR_female_S12 | 16779395 | 2388949 | 5921294 | 1174171 |
| LPR | 9_LPR_female_S2 | 19773472 | 3284957 | 7356944 | 1317193 |
| LPR | 10_LPR_male_S8 | 15172543 | 2379298 | 5457376 | 1070268 |
| LPR | 11_LPR_male_S11 | 14438841 | 2420096 | 5374716 | 1041780 |
| LPR | 12_LPR_male_S1 | 14150680 | 2252137 | 5148394 | 1023886 |

**Table S2.** Assembly statistics for male genomes are shown for the three approaches: SOAPdenovo2 with all reads, SOAPdenovo2 with only reads that did not align to the reference genome, and ABySS. The statistics are scaffold/contig N50, the number of scaffolds/contigs in the assembly, and the total length of the assembled genome. Only scaffolds/contigs that are at least 1 kb long were included in the statistics.

| Assembler | Reads | N50 | Scaffolds/Contigs | Total Length |
| --- | --- | --- | --- | --- |
| SOAPdenovo2 | All | 2491 | 231066 | 522258915 |
| SOAPdenovo2 | MaleUnaligned | 1403 | 4604 | 6769258 |
| ABYSS | All | 1936 | 159804 | 300356116 |

## References

Altschul SF, Madden TL, Schaffer AA, Zhang J, Zhang Z, Miller W, and Lipman DJ. 1997. Gapped BLAST and PSI-BLAST: a new generation of protein database search programs. Nucl. Acids Res. 25: 3389–3402.

Bachtrog D. 2013. Y-chromosome evolution:emerging insights into processes of Y-chromosome degeneration. Nat. Rev. Genet. 14: 113–124.

Bachtrog D, Mank JE, Peichel CL, Kirkpatrick M, Otto SP, Ashman TL, Hahn MW, Kitano J, Mayrose I, Ming R, et al 2014. Sex determination: why so many ways of doing it?PLoS Biol 12: e1001899.

Baker RH, Wilkinson GS. 2010. Comparative genomic hybridization (CGH) reveals a neo-X chromosome and biased gene movement in stalk-eyed flies (genus Teleopsis). PLoS Genet. 6: e1001121.

Benaglia T, Chauveau D, Hunter DR, and Young DS. 2009. mixtools: an R package for analyzing mixture models. J Stat Softw 32: 6.

Beukeboom L and Perrin N. 2014. The Evolution of Sex Determination. Oxford University Press, New York, NY.

Bopp D, Saccone G, and Beye M. 2014. Sex determination in insects: variations on a common theme. Sex. Dev. 8: 20–28.

Boyes JW, Corey MJ, and Paterson HE. 1964. Somatic chromosomes of higher diptera IX. Karyotypes of some muscid species. Can J Cytol 42: 1025–1036.

Bull JJ 1983. Evolution of Sex Determining Mechanisms. Benjamin/Cummings, Menlo Park, CA.

Cakir S and Kence A. 1996. The distribution of males having XY and XX chromosomes in housefly populations (Diptera: Muscidae) of Turkey. Genetica 98: 205–210.

Carvalho AB and Clark AG. 2005. Y chromosome of D. pseudoobscura is not homologous to the ancestral Drosophila Y. Science 307: 108–110.

Carvalho AB and Clark AG. 2013. E cient identi cation of Y chromosome sequences in the human and Drosophila genomes. Genome Res. 23: 1894–1907.

Charlesworth B. 1991. The evolution of sex chromosomes. Science 251: 1030–1033.

Charlesworth B. 1996. The evolution of chromosomal sex determination and dosage compensation. Curr. Biol. 6: 149–162.

Charlesworth B, Coyne JA, and Barton NH. 1987. The relative rates of evolution of sex chromosomes and autosomes. Am. Nat. 130: 113–146.

Charlesworth D, Charlesworth B, and Marais G. 2005. Steps in the evolution of heteromorphic sex chromosomes. Heredity 95: 118–128.

Denholm I, Franco MG, Rubini PG, and Vecchi M. 1983. Identi cation of a male determinant on the X chromosome of housefly (Musca domestica L.) populations in South-East England. Genet. Res. 42: 311–322.

Dobin A, Davis CA, Schlesinger F, Drenkow J, Zaleski C, Jha S, Batut P, Chaisson M, and Gingeras TR. 2013. STAR: ultrafast universal RNA-seq aligner. Bioinformatics 29: 15–21.

Dobzhansky T. 1935. The Y chromosome of Drosophila pseudoobscura. Genetics 20: 366–376.

Dübendorfer A, Hediger M, Burghardt G, and Bopp D. 2002. Musca domestica, a window on the evolution of sex-determining mechanisms in insects. Int J Dev Biol 46: 75–79.

Ellegren H. 2011. Sex-chromosome evolution: recent progress and the inuence of male and female heterogamety. Nat. Rev. Genet. 12: 157–166.

Faber-Hammond JJ, Phillips RB, and Brown KH. 2015. Comparative analysis of the shared sex-determination region (SDR) among salmonid fishes. Genome Biol. Evol. 7: 1972–1987.

Feldmeyer B, Pen I, and Beukeboom LW. 2010. A microsatellite marker linkage map of the housefly, Musca domestica: evidence for male recombination. Insect Mol Biol 19: 575–581.

Filatov DA, Moneger F, Negrutiu I, and Charlesworth D. 2000. Low variability in a Y-linked plant gene and its implications for Y-chromosome evolution. Nature 404: 388–390.

Foster GG, Whitten MJ, Konovalov C, Arnold JTA, and Maffi G. 1981. Autosomal genetic maps of the Australian sheep blowfly, Lucilia cuprina dorsalis R.-D. (Diptera: Calliphoridae), and possible correlations with the linkage maps of Musca domestica L. and Drosophila melanogaster (Mg.). Genet. Res. 37: 55–69.

Graves JAM. 2005. Recycling the Y chromosome. Science 307: 50–51.

Hall AB, Papathanos PA, Sharma A, Cheng C, Akbari OS, Assour L, Bergman NH, Cagnetti A, Crisanti A, Dottorini T, et al 2016. Radical remodeling of the Y chromosome in a recent radiation of malaria mosquitoes. Proc. Natl. Acad. Sci. U.S.A. 113: E2114–E2123.

Hamm RL, Meisel RP, and Scott JG. 2015. The evolving puzzle of autosomal versus Y-linked male determination in Musca domestica. G35: 371–384.

Hamm RL, Shono T, and Scott JG. 2005. A cline in frequency of autosomal males is not associated with insecticide resistance in house fly (Diptera: Muscidae). J. Econ. Entomol. 98: 171–176.

Hediger M, Minet AD, Niessen M, Schmidt R, Hil ker-Kleiner D, Cakir S, Nothiger R, and Dubendorfer A. 1998a. The male-determining activity on the Y chromosome of the housefly (Musca domestica L.) consists of separable elements. Genetics 150: 651–661.

Hediger M, Nfliessen M, Muller-Navia J, Nothiger R, and Dubendorfer A. 1998b. Distribution of heterochromatin on the mitotic chromosomes of Musca domestica L. in relation to the activity of male-determining factors. Chromosoma 107: 267–271.

Hughes JF, Skaletsky H, Koutseva N, Pyntikova T, and Page DC. 2015. Sex chromosome-to-autosome transposition events counter Y-chromosome gene loss in mammals. Genome Biol. 16: 1–9.

Kallman KD. 1984. A new look at sex determination in poeciliid fishes. In Evolutionary Genetics of Fishes (ed.), pp. 95–171. Springer US, Boston, MA.

Koerich LB, Wang X, Clark AG, and Carvalho AB. 2008. Low conservation of gene content in the Drosophila Y chromosome. Nature.

Larracuente AM, Noor MAF, and Clark AG. 2010. Translocation of Y-linked genes to the dot chromosome in Drosophila pseudoobscura. Mol. Biol. Evol. 27: 1612–1620.

Lemos B, Araripe LO, and Hartl DL. 2008. Polymorphic Y chromosomes harbor cryptic variation with manifold functional consequences. Science 319: 91–93.

Leung W, Sha er CD, Cordonnier T, Wong J, Itano MS, Slawson Tempel EE, Kellmann E, Desruisseau DM, Cain C, Carrasquillo R, et al 2010. Evolution of a distinct genomic domain in Drosophila: comparative analysis of the dot chromosome in Drosophila melanogaster and Drosophila virilis. Genetics 185: 1519–1534.

Li H. 2013. Aligning sequence reads, clone sequences and assembly contigs with BWA-MEM. arXiv 1303.3997v2

Li H and Durbin R. 2009. Fast and accurate short read alignment with Burrows-Wheeler transform. Bioinformatics 25: 1754–1760.

Linger RJ, Beliko EJ, and Scott MJ. 2015. Dosage compensation of X-linked Muller element F genes but not X-linked transgenes in the Australian sheep blowfly. PLoS ONE 10: e0141544.

Liu N and Yue X. 2001. Genetics of pyrethroid resistance in a strain (ALHF) of house flies (Diptera: Muscidae). Pestic. Biochem. Physiol. 70: 151–158.

Liu Z, Moore PH, Ma H, Ackerman CM, Ragiba M, Yu Q, Pearl HM, Kim MS, Charlton JW, Stiles JI, et al 2004. A primitive Y chromosome in papaya marks incipient sex chromosome evolution. Nature 427: 348–352.

Love M, Huber W, and Anders S. 2014. Moderated estimation of fold change and dispersion for RNA-seq data with DESeq2. Genome Biol. 15: 550.

Luo R, Liu B, Xie Y, Li Z, Huang W, Yuan J, He G, Chen Y, Pan Q, Liu Y, et al 2012. SOAPdenovo2: an empirically improved memory-efficient short-read de novo assembler. GigaScience 1: 18.

Lyckegaard EM and Clark AG. 1989. Ribosomal DNA and Stellate gene copy number variation on the Y chromosome of Drosophila melanogaster. Proc. Natl. Acad. Sci. U.S.A. 86: 1944–1948.

McKenna A, Hanna M, Banks E, Sivachenko A, Cibulskis K, Kernytsky A, Garimella K, Altshuler D, Gabriel S, Daly M, et al 2010. The Genome Analysis Toolkit: A MapReduce framework for analyzing next-generation DNA sequencing data. Genome Res. 20: 1297–1303.

Meisel RP and Connallon T. 2013. The faster-X effect: integrating theory and data. Trends Genet. 29: 537–544.

Meisel RP, Malone JH, and Clark AG. 2012. Disentangling the relationship between sex-biased gene expression and X-linkage. Genome Res. 22: 1255–1265.

Meisel RP, Scott JG, and Clark AG. 2015. Transcriptome differences between alternative sex determining genotypes in the house fly, Musca domestica. Genome Biol. Evol. 7: 2051–2061.

Miller DD and Roy R. 1964. Further data on Y chromosome types in Drosophila athabasca. Can. J. Genet. Cytol. 6: 334–348.

Miller DD and Stone LE. 1962. Reinvestigation of karyotype in Drosophila a nis Sturtevant and related species. J. Hered. 53: 12–24.

Miura I. 2007. An evolutionary witness: the frog Rana rugosa underwent change of heterogametic sex from XY male to ZW female. Sex. Dev. 1: 323–331.

Muller HJ. 1940. Bearings of the Drosophila’ work on systematics. In The New Systematics (ed. J Huxley), pp.185-268. Clarendon Press, Oxford, Oxford.

Navarro A, Barbadilla A, and Ruiz A. 2000. Effect of inversion polymorphism on the neutral nucleotide variability of linked chromosomal regions in Drosophila. Genetics 155: 685–698.

Patterson JT and Stone WS. 1952. Evolution in the Genus Drosophila. The Macmillan Company, New York.

Rice WR. 1984. Sex chromosomes and the evolution of sexual dimorphism. Evolution 38:735–742.

Rice WR. 1996. Evolution of the Y sex chromosome in animals. Bioscience 46: 331–343.

Roco AS, Olmstead AW, Degitz SJ, Amano T, Zimmerman LB, and Bullejos M. 2015 Coexistence of Y, W, and Z sex chromosomes in Xenopus tropicalis. Proc. Natl. Acad. Sci. U.S.A. 112: 4752–4761.

Ross JA, Urton JR, Boland J, Shapiro MD, and Peichel CL. 2009. Turnover of sex chromosomes in the stickleback fishes (Gasterosteidae). PLoS Genet. 5: e1000391.

Schaeffer SW, Bhutkar A, McAllister BF, Matsuda M, Matzkin LM, O’Grady PM, Rohde C, Valente VLS, Aguade M, Anderson WW, et al 2008. Polytene chromosomal maps of 11 Drosophila species: the order of genomic scaffolds inferred from genetic and physical maps. Genetics 179: 1601–1655.

Scott JG and Georghiou GP. 1985. Rapid development of high-level permethrin resistance in a field-collected strain of the house fly (Diptera: Muscidae) under laboratory selection. J. Econ. Entomol. 78: 316–319.

Scott JG, Sridhar P, and Liu N. 1996. Adult specific expression and induction of cytochrome P 450tpr in house flies. Arch. Insect Biochem. Physiol. 31: 313–323.

Scott JG, Warren WC, Beukeboom LW, Bopp D, Clark AG, Giers SD, Hediger M, Jones AK, Kasai S, Leichter CA, et al 2014. Genome of the house fly, Musca domestica L., a global vector of diseases with adaptations to a septic environment. Genome Biol. 15: 466.

Simpson JT, Wong K, Jackman SD, Schein JE, Jones SJ, and Birol I. 2009. ABySS: a parallel assembler for short read sequence data. Genome Res. 19: 1117–1123.

Smith CD, Shu S, Mungall CJ, and Karpen GH. 2007. The Release 5.1 annotation of Drosophila melanogaster heterochromatin. Science 316: 1586–1591.

Steinemann M and Steinemann S. 1998. Enigma of Y chromosome degeneration: neo-Y and neo-X chromosomes of Drosophila miranda a model for sex chromosome evolution. Genetica 102-103: 409–420.

Stock M, Horn A, Grossen C, Lindtke D, Sermier R, Betto-Colliard C, Dufresnes C, Bonjour E, Dumas Z, Luquet E, et al 2011. Ever-young sex chromosomes in european tree frogs. PLoS Biol. 9: e1001062.

Sturgill D, Zhang Y, Parisi M, and Oliver B. 2007. Demasculinization of X chromosomes in the Drosophila genus. Nature 450: 238–241.

Tomita T and Wada Y. 1989. Multifactorial sex determination in natural populations of the housefly (Musca domestica) in Japan. Jpn. J. Genet. 64: 373–382.

Traut W and Willhoeft U. 1990. A jumping sex determining factor in the fly Megaselia scalaris. Chromosoma 99: 407–412.

Vallender EJ and Lahn BT. 2006. Multiple independent origins of sex chromosomes in amniotes. Proc. Natl. Acad. Sci. U.S.A. 103: 18031–18032.

Veyrunes F, Catalan J, Sicard B, Robinson TJ, Duplantier JM, Granjon L, Dobigny G, and Britton-Davidian J. 2004. Autosome and sex chromosome diversity among the African pygmy mice, subgenus Nannomys (Murinae; Mus). Chromosome Res. 12: 369–382.

Vicoso B and Bachtrog D. 2013. Reversal of an ancient sex chromosome to an autosome in Drosophila. Nature 499: 332–335.

Vicoso B and Bachtrog D. 2015. Numerous transitions of sex chromosomes in Diptera. PLoS Biol. 13: e1002078.

Vicoso B and Charlesworth B. 2006. Evolution on the X chromosome: unusual patterns and processes. Nat. Rev. Genet. 7: 645–653.

Vicoso B, Kaiser VB, and Bachtrog D. 2013. Sex-biased gene expression at homomorphic sex chromosomes in emus and its implication for sex chromosome evolution. Proc. Natl. Acad. Sci. U.S.A. 110: 6453–6458.

Wagoner DE. 1967. Linkage group-karyotype correlation in the house fly determined by cytological analysis of X-ray induced translocations. Genetics 57: 729–739.

Wei KHC, Grenier JK, Barbash DA, and Clark AG. 2014. Correlated variation and population differentiation in satellite DNA abundance among lines of Drosophila melanogaster. Proc. Natl. Acad. Sci. U.S.A. 111: 18793–18798.

Weller GL and Foster GG. 1993. Genetic maps of the sheep blowfly Lucilia cuprina: linkage-group correlations with other dipteran genera. Genome 36: 495–506.

Woram RA, Gharbi K, Sakamoto T, Hoyheim B, Holm LE, Naish K, McGowan C, Ferguson MM, Phillips RB, Stein J, et al2003. Comparative genome analysis of the primary sex-determining locus in salmonid fishes. Genome Res. 13: 272–280.

Yazdi HP and Ellegren H. 2014. Old but not (so) degenerated|slow evolution of largely homomorphic sex chromosomes in ratites. Molecular Biology and Evolution 31: 1444–1453.

